# Cardiac Specific Overexpression of Adenylyl Cyclase 8 Reprograms the Mouse Sinoatrial Node Transcriptome

**DOI:** 10.1101/2023.09.26.559627

**Authors:** Jia-Hua Qu, Kirill V. Tarasov, Khalid Chakir, Edward G. Lakatta

## Abstract

A coupled-clock system intrinsic to sinoatrial node (SAN) pacemaker cells that regulates the rate and rhythm of spontaneous action potential firing, is activated by Ca^2+^/calmodulin-stimulated Adenylyl Cyclase (AC) types 8 and 1. Our previous work in mice with cardiac specific overexpression of human AC8 gene (TGAC8) discovered that compared to its wild-type (WT) littermates, the heart rate (HR) of TGAC8 is elevated (by about 30%, 24 hours a day, 7days a week), and that the TGAC8 heart rhythm is markedly coherent, i.e., the HR Variability (HRV) lacks complexity, similar to that associated with aging or cardiac pathology. Reprogramming of molecular mechanisms, particularly SAN transcriptomic regulation that underlies the remarkable chronic shift in HR and HRV, however, has not been delineated. We conducted deep RNA sequencing (RNA-seq) in TGAC8 and WT SANs, using Mm10plus with human ADCY8 DNA sequence as the reference genome. Utilizing multiple bioinformatic techniques, we not only profiled the expression of marker genes related to SAN functions and AC-cAMP-PKA signaling, but also discovered negatively enriched hub pathways that differed in TGAC8 vs. WT, the top three being OXPHOS, ribosome, and cardiac muscle contraction. In contrast, signaling pathways related to inositol phosphate and its metabolism were positively enriched in TGAC8. Further, we identified two transcription regulators, KDM5A and PPARGC1A, that mediate effects of TGAC8 on ribosome and mitochondria. In summary, reprogrammed transcriptional regulation increases the HR of TGAC8 at the cost of impairment of other SAN cell functions, i.e., altered ribosome and inositol phosphate signaling and its crosstalk with mitochondria. Because these cell signaling alterations are associated with cardiac aging and age-associated CVDs, the TGAC8 mouse appears to be an ideal model in which to probe for potential therapeutic targets for cardiac aging and age-associated CVDs.

## Introduction

According to the report from World Health Organization (https://www.who.int/cardiovascular_diseases/about_cvd/en/), cardiovascular diseases (CVDs) are the leading cause of death globally. The decline of cardiac performance in CVDs may result from an impaired contractile function^1,2^. As one of the most important regulators of contraction, cAMP is catalyzed by adenylyl cyclases (ACs) upon activation by catecholamines binding to β-adrenergic receptors (β-ARs), and degraded by phosphodiesterases (PDEs)^3^. The activated ACs-cAMP-PKA signaling pathway promotes the phosphorylation of key proteins, contributing to strong positive inotropic, lusitropic, and chronotropic responses^3^. Therefore, the desensitization of the β-adrenergic signaling pathway, with the down-regulation of β1-ARs and ACs, disrupts the excitation-contraction coupling, resulting in CVDs^4^.

Contrary to two main AC isoforms expressed in the heart, AC5 and AC6, two additional isoforms in sinoatrial node (SAN), AC1 and AC8, are activated by Ca^2+^/calmodulin and insensitive to subunit of heterotrimeric G proteins (Gαs) in vitro^5,6^. In pacemaker cells, the AC8-cAMP-PKA signaling pathway induces the phosphorylation of Ca^2+^ cycling proteins to generate the action potential^5^. In our previous work using the mouse model with cardiac specific overexpression of human AC8 gene (TGAC8), we have discovered that the heart rate (HR) of TGAC8 is increased approximately 30% compared with the wild-type (WT) littermate, and the intrinsic coupled-clock system (in the absence of β1-ARs stimulation) within the SAN cells is crucially dependent on the activation of AC8, but the overall pattern of marked coherency or loss of complexity (HR variability) within the TGAC8 heart rhythm is similar to that associated with aging or cardiac pathology^7^. Although we have already profiled the changes in the TGAC8 mouse left ventricle (LV) at RNA, protein and phosphorylation levels^8,9^, and have characterized the various cellular functions at single nucleus resolution in C57/BL6 mouse SAN^10^, the molecular mechanisms, especially the transcriptomic regulation in SAN, by which TGAC8 regulates SAN functions and its long-term effect, remain elusive.

To unravel the underlying molecular mechanisms, in the present research, we conducted deep RNA sequencing (RNA-seq) to profile the SAN transcriptome in mouse (TGAC8 vs. WT). Utilizing multiple bioinformatic techniques, we enriched representative pathways and discovered an unexpected downregulation of components and functions of mitochondria and ribosome, which may be associated with CVDs in long term of chronic activation of AC8 in the heart. Our results provide valuable transcriptomic data for further research on the regulation in cardiac function, especially through the ACs-cAMP-PKA signaling pathway.

## Materials and Methods

### 1. Animals

The cardiac-specific overexpression of AC8 (TGAC8) mouse, generated by inserting a cDNA coding for human AC8 after the murine a-myosin heavy chain promoter^6^, was a gift from Nicole Defer/Jacques Hanoune, Unite de Recherches, INSERM U-99, Hôpital Henri Mondor, F-94010 Créteil, France. Twelve-week old male TGAC mice were used as study cases, and age matched WT littermates, in the C57/BL6 background, were used as controls. All studies were performed in accordance with the Guide for the Care and Use of Laboratory Animals published by the National Institutes of Health (NIH Publication no. 85-23, revised 1996). The experimental protocols were approved by the Animal Care and Use Committee of the National Institutes of Health (protocol #441-LCS-2016).

### 2. Sinoatrial node (SAN) and SAN cell isolation

SAN and SAN cells were isolated in the same way described in our previous paper^7^. Briefly, after anesthetization of TGAC8 and WT mice, the hearts were quickly removed and SANs were dissected between inferior and superior vena cava, crista terminalis, and intra-atrial septum and cut into strips perpendicular to the crista terminalis. The isolated SANs were washed (X3) in the modified low Ca^2+^ Tyrode solution before enzymatic digestion. The digested SANs were washed (X3) in modified high potassium solution before dispersion by gentle pipetting. The dispersed SAN cells were stored at 4℃ for following experiments.

### 3. RNA extraction, cDNA library preparation, and RNA-seq

By company. We sent eight TGAC8 and eight WT mouse SANs to DNA Link USA company to conduct total RNA-seq. RNAs were extracted from each SNAs separately and used as template for cDNA library preparation following the instruction manual of SMARTer Stranded Total RNA-Seq Kit-Pico/Clontech Laboratories, Lnc. The sequencing was conducted on the Illumina NextSeq 500 platform and the data were obtained within the basecaling software RTA2.0.

### 4. RNA-seq data processing

The raw reads were trimmed within Cutadapt and aligned within SAMtools. Using Mm10plus and ADCY8 as the reference genome, the aligned reads were mapped to calculate the raw counts. After quality control and filtration, we selected six WT samples (WS1, WS2, WS3, WS4, WS5 and WS 8) and five TGAC8 samples (MS1, MS2, MS4, MS5 and MS8) for the subsequent differential expression gene (DEG) analysis using DESeq2 package in R language, and identified 16575 genes. The normalized counts by DESeq2 were imported into Partek Flow to perform PCA and generate expression heat map. Furtherly, we calculated sample distances using the Euclidean method for rows and columns distance clustering and generated matrix using the ward.D method for matrix clustering in R language. The adjusted p-value and fold change calculated by DESeq2 were used to draw volcano plot to figure out the expression changes of TGAC8 SAN transcriptome. Specifically, 352 genes were downregulated (-log10(adjusted p-value) > 1.3 and log2(fold change) < 0) and 323 upregulated ((-log10(adjusted p-value) > 1.3 and log2(fold change) > 0).

### 5. Marker genes selection and analysis

The expressions of several families of genes encoding proteins related to ion channels and AC8-cAMP-PKA pathway, i.e., sodium/calcium exchanger in solute carrier family (Slc), hyperpolarization activated cyclic nucleotide gated potassium channel (Hcn), S100 calcium binding protein (S100), sodium voltage-gated channel (Na+), potassium voltage-gated channel (K+), calcium voltage-gated channel (Ca2+), stromal interaction molecule (Stim), phosphodiesterase (Pde), protein kinase CAMP-dependent regulation proteins (PKA), and sarcolipin (Sln), were extracted from TGAC8 SAN RNA-seq data and visualized in heat map. The significant changes of these genes were further displayed in volcano plot and bar plot. All analyses were performed and all plots were produced in R language.

### 6. Ingenuity pathway analysis (IPA)

All identified genes were uploaded to IPA software (QIAGEN, March 2020)^11^. The Ingenuity knowledge base (gene only) about all species and all tissues was selected as the reference set. All Ingenuity supported third party information and Ingenuity expert information were adopted as the data sources, and only experimentally observed relationships were selected to increase confidence. Setting the cutoff as adjusted p-value < 0.05, IPA recognized 636 differential expressed genes, including 321 down and 315 up, which were used for subsequent canonical pathway analysis, upstream regulator prediction, disease state analysis match, and regulatory pathway construction. Specifically, OXPHOS pathway and apelin cardiomyocyte signaling pathway were visualized, and the regulatory pathway starting from KDM5A and PPARGC1A was generated in IPA.

### 7. KEGG pathway analysis using pathfindR package

To reduce the redundancy and find the most important pathways, KEGG pathway analysis was performed using pathfindR package in R language^12^. The KEGG mouse (mmu) database was used as background gene set. Besides normal KEGG enrichment, we generated term-gene graph to cluster all pathways into seven clusters and select the hub pathway in each cluster. We also displayed the pathway level heat map for individual sample to clarify the contribution of each sample to the pathway enrichment.

### 8. Gene set enrichment analysis (GSEA)

To take all identified genes into consideration and preclude bias due to arbitrary cutoff, we uploaded all identified genes into GSEA software^13^ for analyses within four databases, KEGG, GO_BP, GO_CC, and GO_MF. The software can recognize 16571 native features. After collapsing features into gene symbols, there are 12873 genes remained. The calculation parameters in GSEA were as below: (1) gene-set permutation; (2) permutation number: 1000; (3) scoring_scheme: weighted; (4) norm: meandiv; (5) mode: Max_probe; (6) include_only_symbols: true; (7) set_max: 500; (8) set_min: 15; (9) metric: Signal2Noise; (10) order: descending. The top 20 significantly enriched pathways sorted by normalized enrichment score (NES) within each database were used for generation of bubble graphs and representative GSEA enrichment plots.

### 9. RNA extraction, cDNA synthesis, qRT-PCR, and data analysis

The preparation for and performance of qRT-PCR were done as stated in our previous paper in detail^7^. Briefly, RNAs were extracted from isolated mouse SAN with RNeasy Mini Kit (Qiagen, Valencia, CA, United States) and DNAse. 2 mg of total RNA was used for cDNA synthesis with MMLV reverse transcriptase (Promega) in final 50 mL volume. Primers used for each transcript assessed are listed in Supplementary Table (in preparation). qRT-PCR was performed on QuantStudio 6 Flex Real-Time PCR System (Thermo Fisher Scientific) with 384-well platform. Reaction was performed with FastStart Universal SYBR Green Master Kit with Rox (Roche). Normalized to expression of HPRT level, qRT-PCR analysis was performed using ddCt method.

### 10. Statistical analyses

Transcriptomic data were processed in Partek^®^ Flow^®^ software, version 9.0 Copyright ©; 2020 Partek Inc., St. Louis, MO, USA, by utilizing the computational resources of the NIH HPC Biowulf cluster (http://hpc.nih.gov). Most statistical analyses were performed using RStudio (version: 1.1.463) in R language (version: 3.5.3). Some other data processing and statistics were conducted in GraphPad Prism (version: 7) and Microsoft Excel (version: 2019). Adobe Illustrator (version: CC 2019) were also used for graphics without changing data trend or structure. For qRT-PCR analysis, the two-sided unpaired Student’s t-test was used to compare the mRNA expression between TGAC8 and WT SAN, and p-value < 0.05 was taken as statistically significant.

## Results

### 1. RNA sequencing on mouse SAN tissue

To study the molecular mechanism, we isolated SANs from twelve-week-old male TGAC8 mice and their WT littermates to conduct deep RNA sequencing (RNA-seq). TG and WT samples were separated into two groups in primary components analysis (PCA) (Figure A1). Furtherly, we calculated sample distances and observed that the inner group sample distance was smaller than inter group sample distance (Figure 1B). We also scaled and visualized the expressions of all identified genes in heatmap, and observed two hierarchical clusters, indicating the differential regulation between the two groups of samples (Figure 1C). Setting the cutoff as -log10(adjusted p-value) > 1.3, i.e., adjusted p-value < 0.05, we obtained 675 differentially regulated (up and down) genes, among which 352 genes were downregulated (down) and 323 upregulated (up) (Figure 1D).

**Figure 1.**
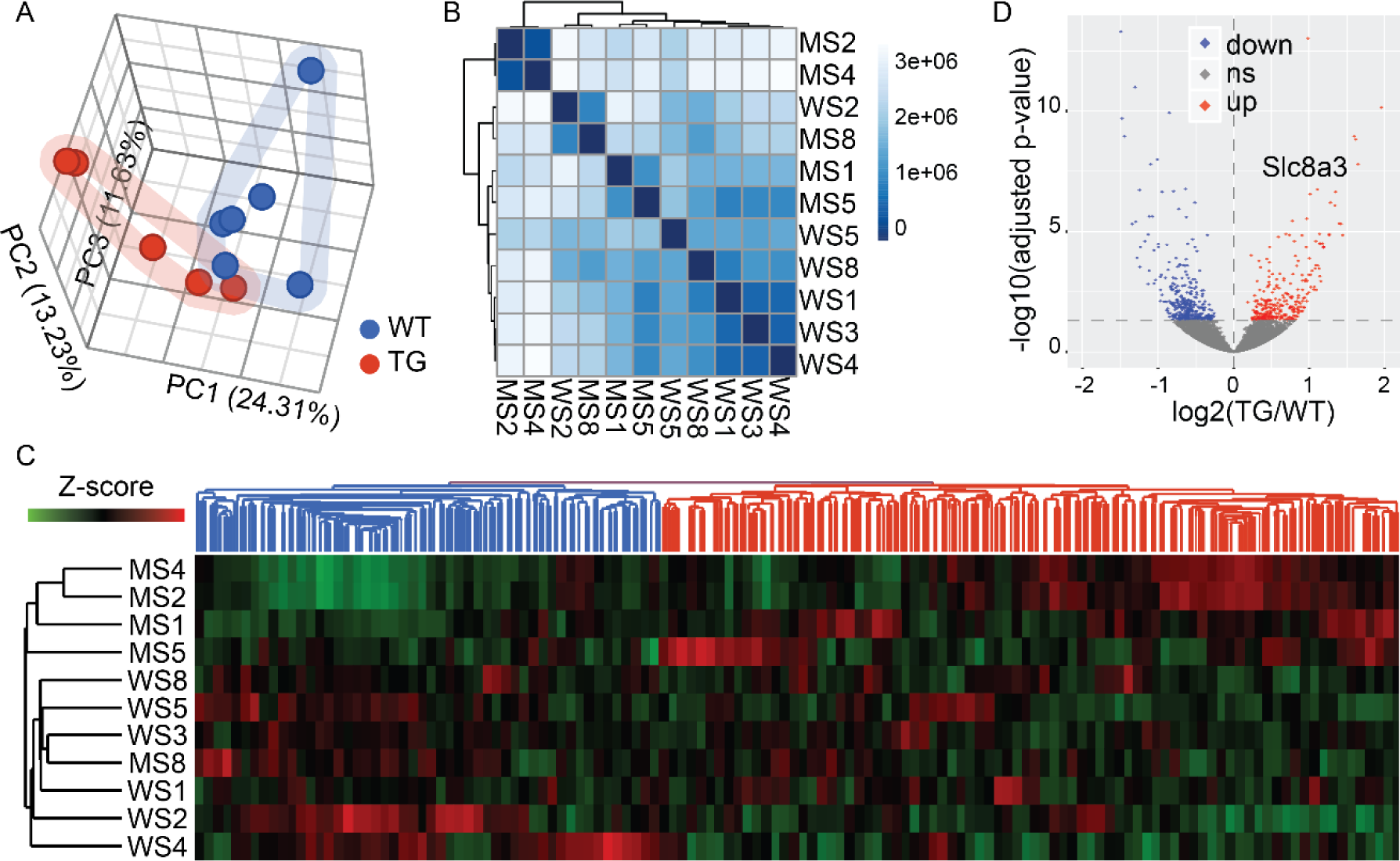
Conduction of RNA sequencing on samples of TGAC8 vs. WT mouse SAN. **A-C**. PCA (A), Sample distance matrix (B), and hierarchial cluster (C) of all samples used in the TGAC8 SAN transcriptome (WT: WS1, WS2, WS3, WS4, WS5, WS8; TGAC8: MS1, MS2, MS4, MS5, MS8). **D**. Volcano plot of TGAC8 SAN RNA-seq data. Genes with adjusted p-value less than 0.05 are regarded as significantly regulated by the overexpression of AC8, among which genes with log2(TG/WT) more than zero and less than zero are marked as red (up) and blue dots (down) respectively. Other genes are nonsignificantly regulated and marked as grey dots (ns). One of the most significantly upregulated genes, Slc8a3, is labeled besides its dot.

As SAN is the main pacemaker of the heart and AC8-cAMP-PKA axis is pivotal to excitation-contraction coupling, we profiled the expression of several families of genes encoding proteins related to ion channels and AC8-cAMP-PKA pathway, i.e., sodium/calcium exchanger in solute carrier family (Slc), hyperpolarization activated cyclic nucleotide gated potassium channel (Hcn), S100 calcium binding protein (S100), sodium voltage-gated channel (Na^+^), potassium voltage-gated channel (K^+^), calcium voltage-gated channel (Ca^2+^), stromal interaction molecule (Stim), phosphodiesterase (Pde), protein kinase cAMP-dependent regulation proteins (PKA), and sarcolipin (Sln) (Figure 2A), among which there were 13 genes regulated significantly by overexpression of AC8 (Figure 2B). Particularly, one member in solute carrier family, Slc8a3, was increased most significantly (Figure 1D and Figure 2C), and two members in phosphodiesterase were also upregulated as feedback to the overexpression of AC8 to finetune the content of cAMP inside SAN (Figure 2C). The characterization of RNA-seq data demonstrated that the cardiac-specific overexpression of AC8 reprogramed the transcriptome in SAN and regulated expression of genes related to cardiac functions.

**Figure 2.**
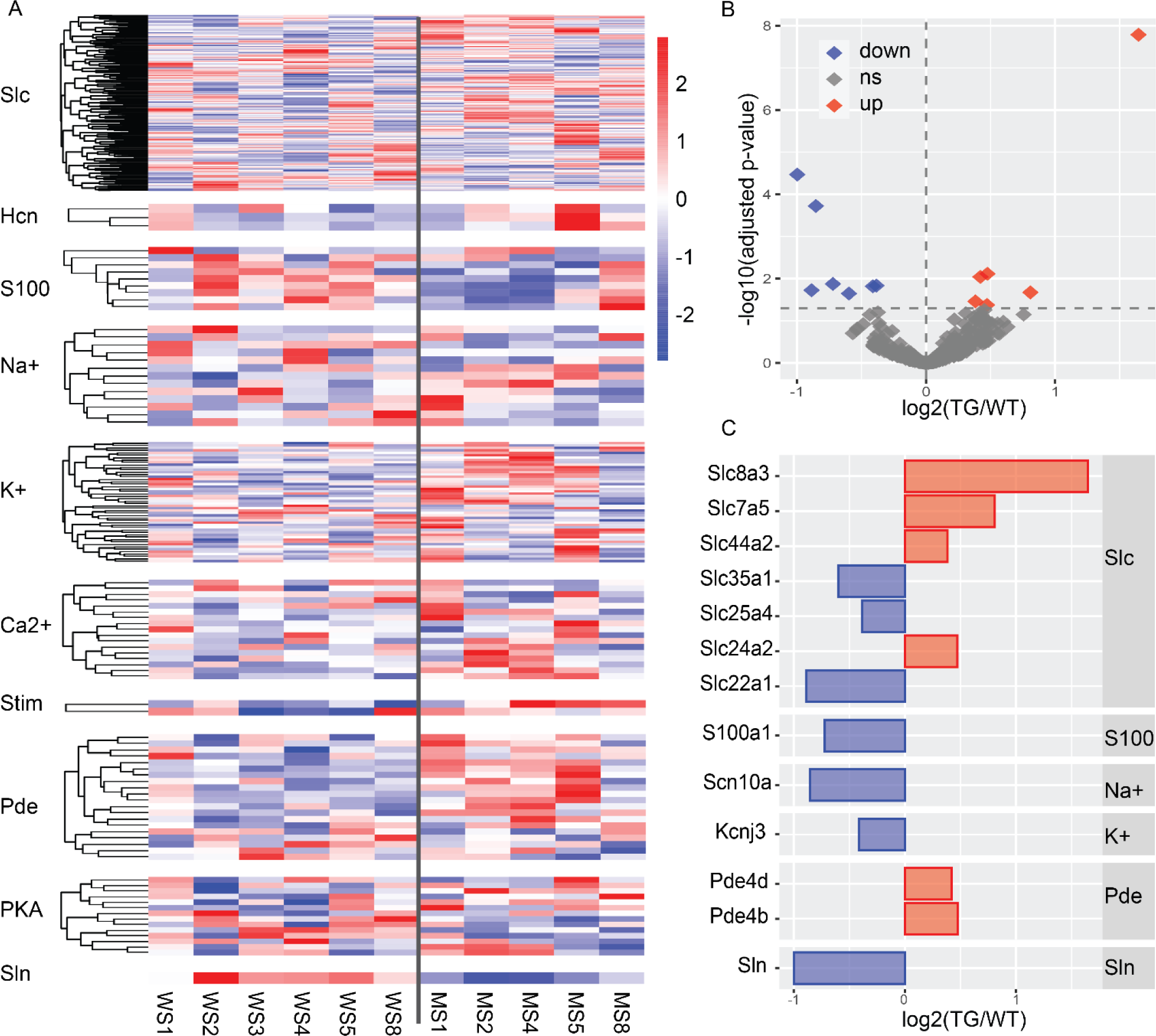
Marker genes in TGAC8 SAN transcriptome. **A**. Heatmap of selected families of genes related to ion channels and AC8-cAMP-PKA pathway. **B**. Volcano plot of all selected genes in A. Red dots represent upregulated genes (adjusted p-value < 0.05 and log2FoldChange > 0); blue dots represent downregulated genes (adjusted p-value < 0.05 and log2FoldChange < 0); grey dots represent nonsignificantly regulated genes (adjusted p-value ≥ 0.05). **C**. Bar plot of expressions of genes that are significantly regulated in AC8 SAN (up and down).

### 2. Functional enrichment of TGAC8 SAN transcriptome

To reveal the functions of the reprogramed SAN transcriptome, we performed functional analysis using Ingenuity Pathway Analysis (IPA) software. The top 25 enriched canonical pathways sharing same genes interacted with each other. As expected, protein kinase A signaling was in the center of the aggregated pathways (Figure 3A), confirming the activation of AC8-cAMP-PKA signaling pathway in our transgenic mouse model (TGAC8). With enrichment p-value and activation prediction z-score, pathways related to mitochondrial functions were unexpectedly enriched and ranked at the top, sorted by - log10(p-value) (Figure 3B).

**Figure 3.**
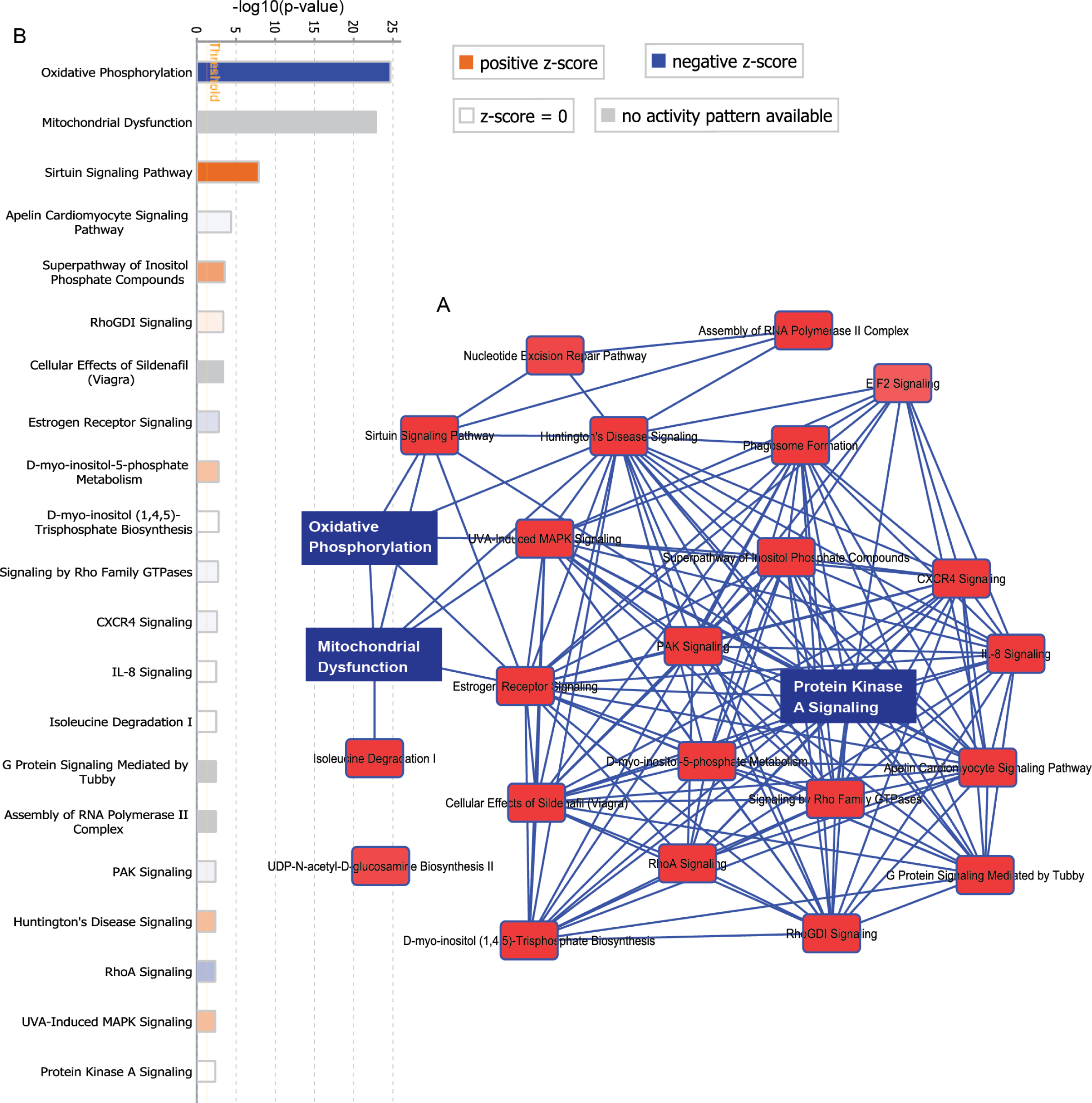
Ingenuity Pathway Analysis of TGAC8 SAN transcriptome. **A**. Overlapping network of top 25 pathways. Three representative pathways, Oxidative Phosphorylation, Mitochondrial Dysfunction, and Protein Kinase A Signaling, are marked in blue rectangles. **B**. Top 21 canonical pathways enriched in IPA, ranked by -log10(p-value). The 20^th^ and 21^st^ pathways have the same -log10(p-value). Bar color indicates activation state based on the uploaded transcriptome into IPA. Orange bars with positive z-scores and blue bars with negative z-scores denote activated and inhibited pathways respectively. The darker the more significant. White bars denote pathways insignificantly activated or inhibited with z-score equal to zero. Grey bars denote pathways for which no activity pattern is available within IPA knowledgebase.

We also conducted KEGG pathway analysis using pathfindR package in R language to study the functions of TGAC8 SAN transcriptome from another respect. Similar to results from IPA, oxidative phosphorylation (OXPHOS) pathway was also enriched and ranked as the top pathway among all enriched KEGG pathways (Figure 4A). Many genes were shared by multiple pathways (Figure 4B-4C), so to reduce the redundancy and find the most important molecular pathways, we clustered the enriched pathways into seven clusters, from which we selected one hub pathway as the representative for each pathway cluster (Figure 4D). The OXPHOS pathway was still the most significant pathway as the hub in cluster 1. In addition, ribosome and cardiac muscle contraction were also enriched and served as the hub in cluster 2 and cluster 3 respectively (Figure 4D). Interestingly, from the term-gene graph and pathway heatmap, we observed that the majority of associated genes were downregulated, especially genes in the top three hub pathways, i.e., OXPHOS, ribosome, and cardiac muscle contraction (Figure 4B-4C, 4E). Therefore, the regulation of these three pathways may be the molecular mechanism underlying the TGAC8 SAN transcriptome, and we continued to analyze them in depth.

**Figure 4.**
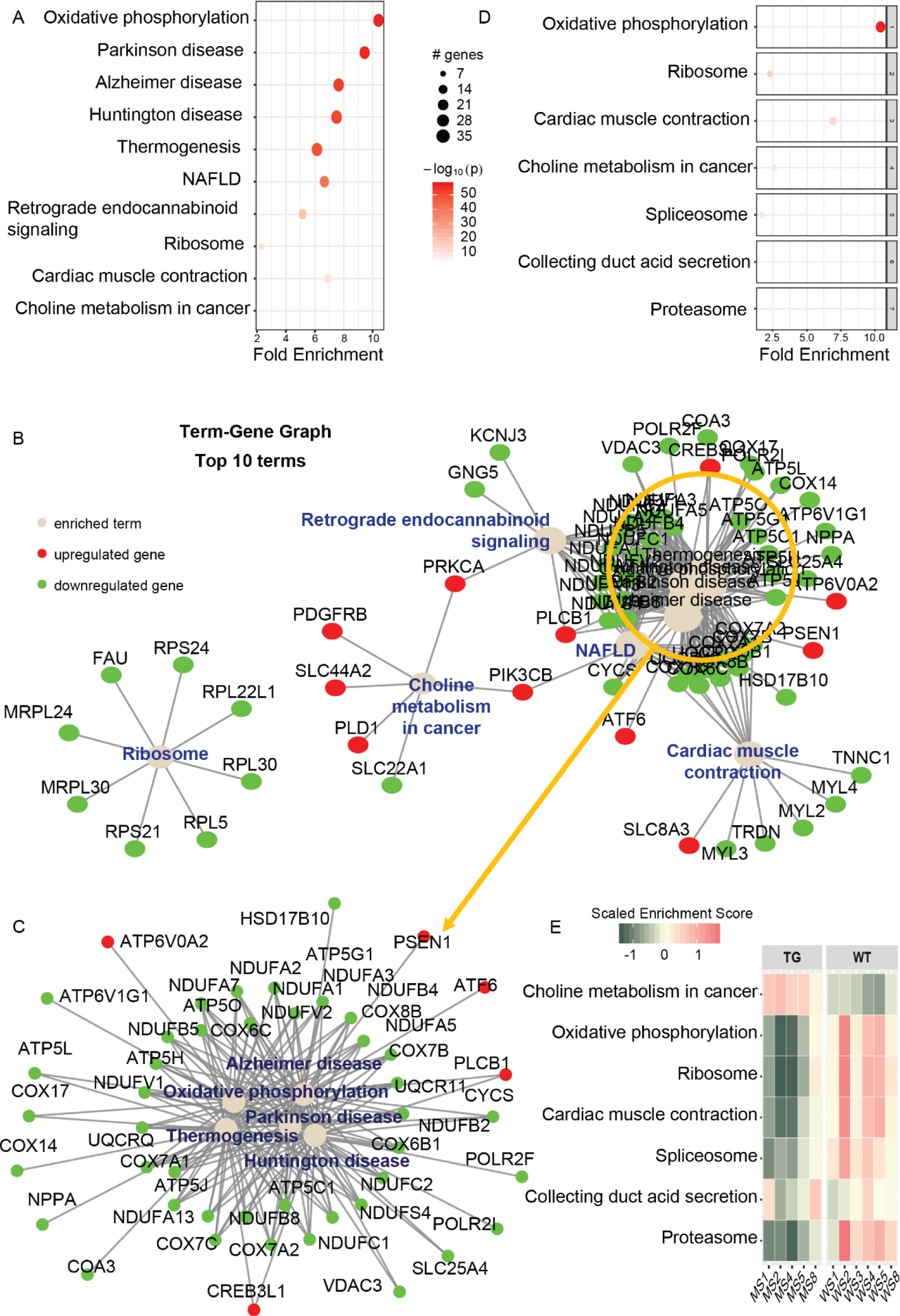
KEGG pathway clustering analysis of TGAC8 SAN transcriptome. **A**. Bubble graph of KEGG pathway analysis of TGAC8 SAN transcriptome. X-axis is fold enrichment of pathway, bubble size represents number of genes in a pathway, and bubble color gradient indicates significance with -log10(p-value). **B**. Term-gene graph of top 10 terms, ranked by fold enrichment. Grey dot denotes enriched term. Red and green dots denote upregulated and downregulated genes respectively. Some genes are enriched in more than one pathway. The pathways on the top right are aggregated and thus zoomed in as Figure **C**. **D**. Bubble graph of hub KEGG pathways in each pathway cluster using pathfindR package. Legend same to Figure A. **E**. Heat map of term score in each sample. Color gradient indicates scaled enrichment score. The more red, the more enriched.

### 3. Suppressed mitochondria and mitochondrial function pathways

The fitness of SAN relies on normal mitochondrial functions to generate enough energy and maintain energy homeostasis. The above analyses in IPA and KEGG, with the cutoff p-value < 0.05, had already implied the inhibition of mitochondrial function in TGAC8 mouse. To take all identified genes into consideration, we made use of gene set enrichment analysis (GSEA). In congruent with the above analyses, many mitochondrial components and mitochondrial functions were enriched without the requirement of any cutoff of p-value or fold change in GSEA (Figure 5A). Regardless of the cutoff, more genes in the gene set of OXPHOS were used for calculation and most genes were still enriched as downregulated in TGAC8 mouse, as shown in the representative GSEA enrichment plots (Figure 5B). Therefore, multiple functional enrichment methods, including IPA, pathfindR, and GSEA, drew the same conclusion: suppressed mitochondria and mitochondrial function in TGAC8 vs. WT.

**Figure 5.**
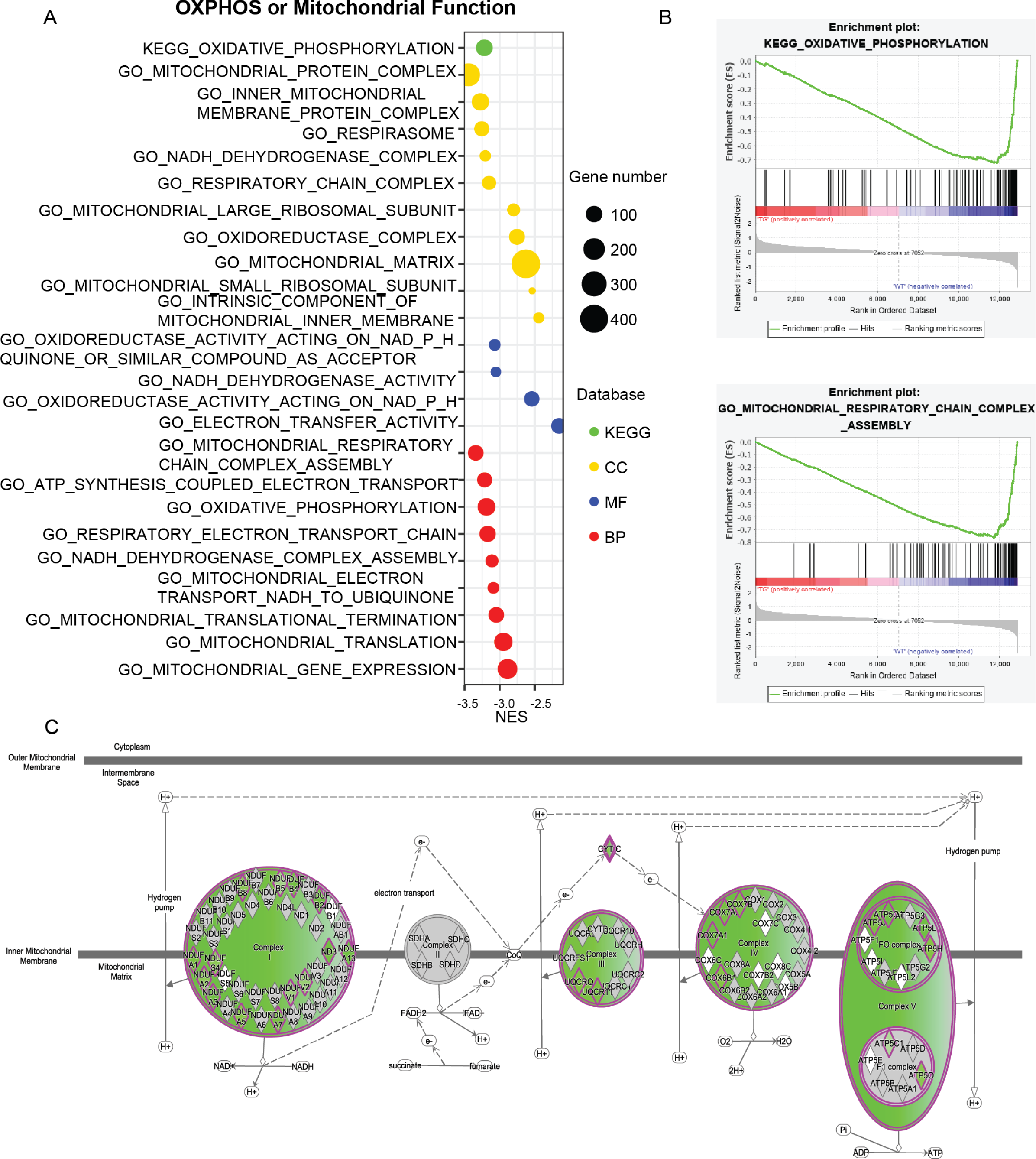
Downregulated mitochondria and mitochondrial functions in TGAC8 SAN. **A**. Bubble graph of GSEA of TGAC8 SAN transcriptome. X-axis is normalized enrichment score (NES) in GSEA, bubble size represents number of genes in a term, and bubble color indicates enrichment using four different databases (KEGG, GO_CC, GO_MF, GO_BP). **B**. Two representative GSEA enrichment plots. Top: enriched using KEGG database; bottom: GO_BP. **C**. Visualization of changed genes in the term of oxidative phosphorylation, generated in IPA. Genes with purple outline are changed significantly. All genes are in green, indicating decreased.

To visualize individual differentially regulated gene related to mitochondria and mitochondrial function, we visualized the OXPHOS pathway within IPA. Significantly regulated genes (adjusted p-value < 0.05) encode subunits in OXPHOS complex I, III, IV and V (colored with purple outlines), and all of them were downregulated (filled in green) (Figure 5C).

To confirm the RNA-seq results, we would perform q-RT PCR to evaluate the expression of the transcribed mRNAs. Mitochondrial function was also analyzed via WB, activity kits, and seahorse analysis, especially for complex I, III, IV, and V.

### 4. Suppressed ribosome and translation pathways

The translation process is one stage of the cell protein quality control, which consumes large amount of energy produced in mitochondria. Mitochondria, as a semi-autonomic organelle, consists of proteins that are synthesized in cytoplasm and mitochondria itself. The majority of mitochondrial proteins are translated by cytosolic ribosome, and a small portion is translated by mitochondrial ribosome. The enrichment of ribosome as the hub in cluster 2 of KEGG pathways (Figure 4D) prompted us to further analyze ribosome and translation. In agreement with the enrichments of both cytosolic ribosome and mitochondrial ribosome subunits, translation inside and outside of mitochondria were also enriched in GSEA (Figure 6A). In line with the downregulation of genes related to ribosome that were displayed in term-gene graph (Figure 3B), the peaks in GSEA enrichment plots of ribosome and translation were downward, indicating the suppression of translation (Figure 6B). Particularly, the inhibited mitochondrial ribosome and mitochondrial translation implied the possible reciprocal regulation between mitochondria and ribosome: suppressed mitochondria produce less energy for translation, while downregulated ribosome and weakened translation generate fewer mitochondrial proteins.

**Figure 6.**
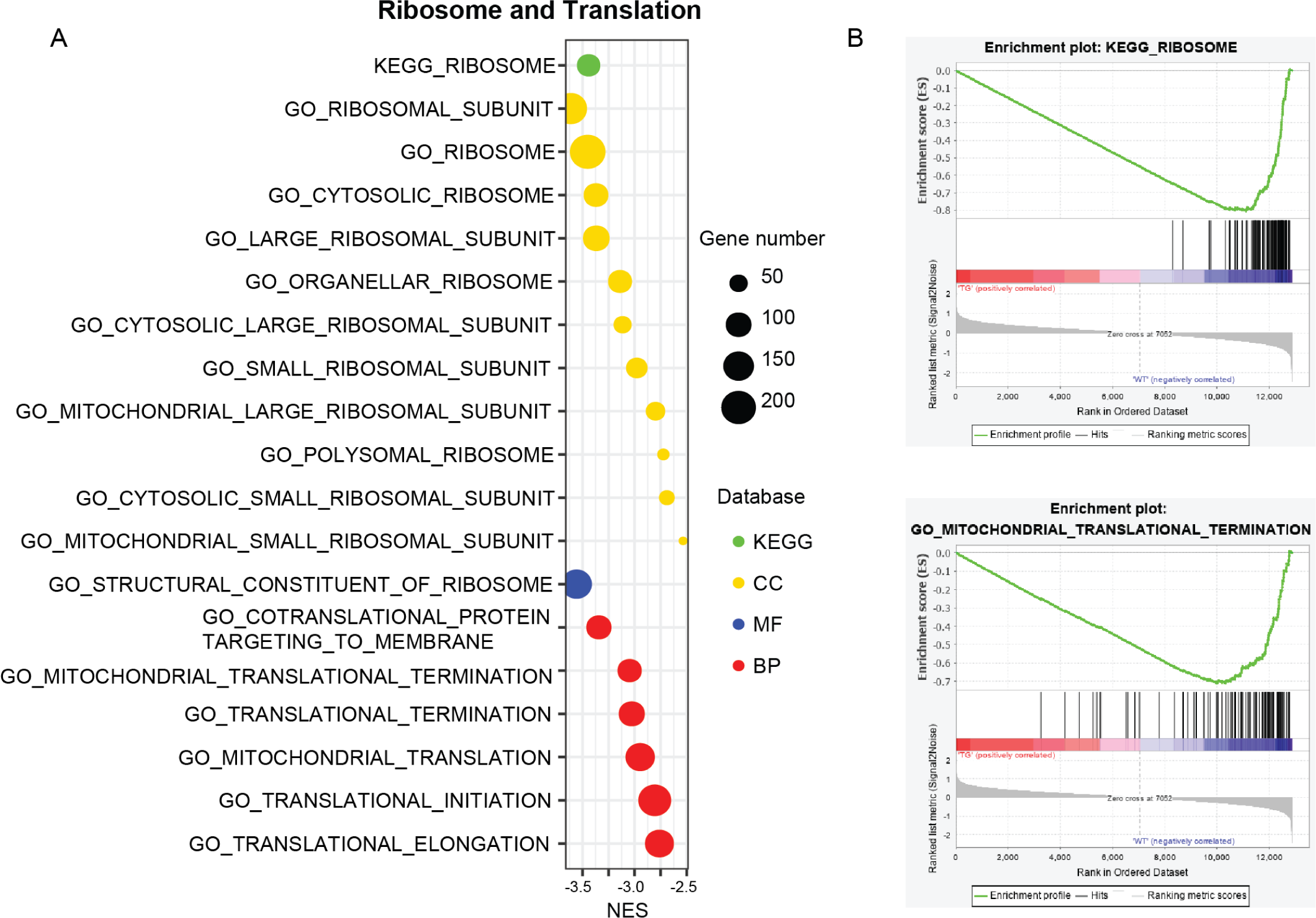
Downregulated ribosome and translation in TGAC8 SAN. **A**. Bubble graph of GSEA of TGAC8 SAN transcriptome. X-axis is normalized enrichment score (NES) in GSEA, bubble size represents number of genes in a term, and bubble color indicates enrichment using four different databases (KEGG, GO_CC, GO_MF, GO_BP). **B**. Two representative GSEA enrichment plots. Top: enriched using KEGG database; bottom: GO_BP.

We would perform qRT-PCR to confirm the downregulated transcription of ribosomal components and do protein synthesis detection and WB to verify the suppressed translation.

### 5. Association with impaired cardiac functions

Intriguingly, only inositol phosphate related pathways were significantly positively enriched within GSEA (Figure 7). As a family of calcium channels, inositol 1,4,5-trisphosphate receptors (IP3Rs) are ubiquitously expressed in all tissues^8^. In the heart, IP3Rs have been associated with regulation of cardiomyocyte function ranging from the regulation of pacemaking, excitation–contraction and excitation–transcription coupling^14,15^. The upregulated inositol phosphate increases the inner cellular Ca^2+^ to enhance cardiac contraction^14,15^, which perhaps mediates the effect of overexpression of AC8 to increase the HR^7^. However, the abnormally activated IP3Rs signaling may contribute to CVDs, like arrhythmias, hypertrophy and heart failure^14,15^. Mechanistically, inositol 1,4,5-trisphosphate (IP3) is involved in the flow of Ca^2+^ between mitochondria and endoplasmic/sarcoplasmic reticulum^16^. In cardiomyocytes, IP3 regulates mitochondrial Ca^2+^ uptake through the mitochondrial ryanodine receptor (RyR) to influence adenosine triphosphate (ATP) production, which may be one mechanism how abnormal inositol phosphate metabolism impairs the cardiac function^17^.

**Figure 7.**
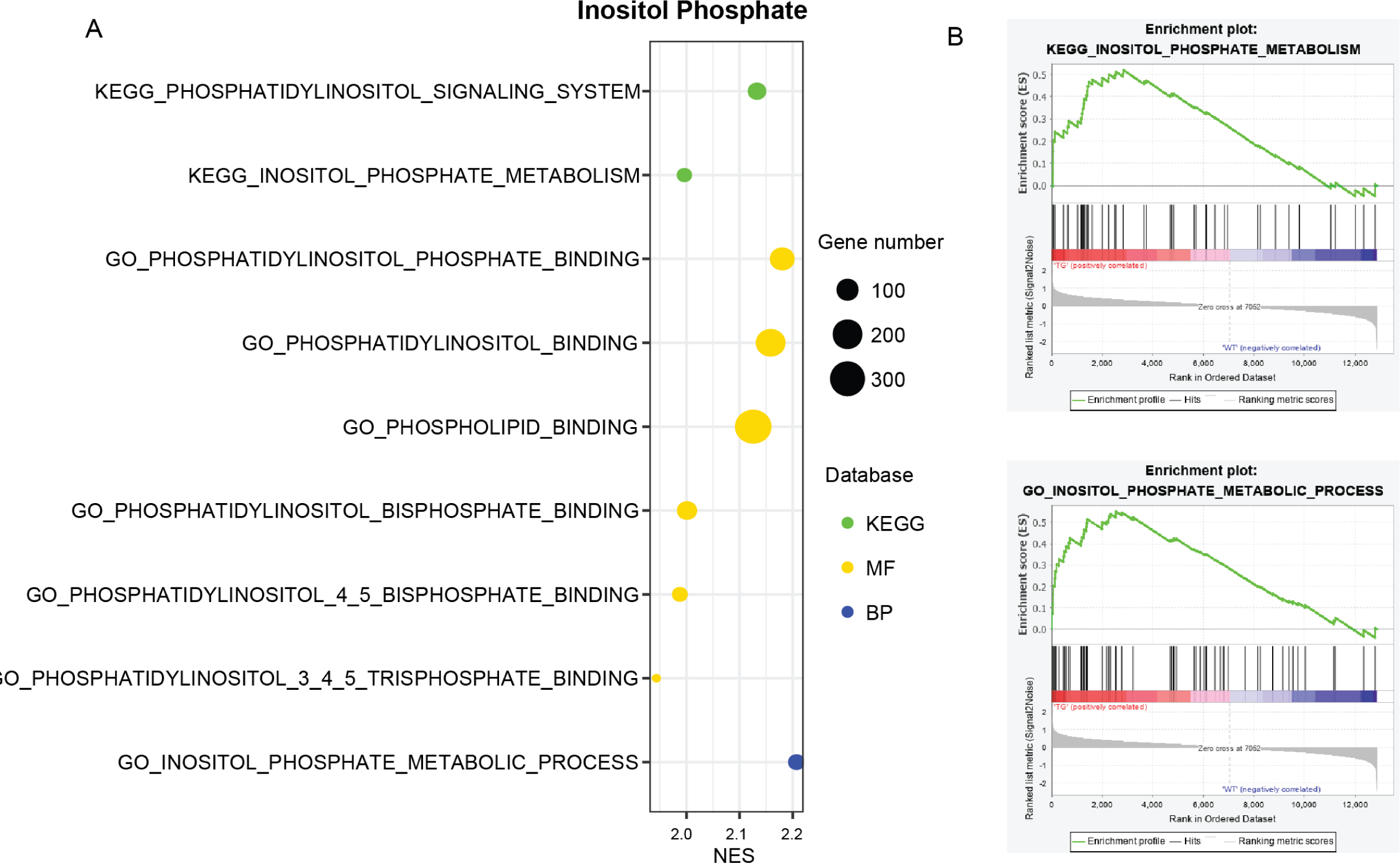
Upregulated inositol phosphate and its metabolism in TGAC8 SAN. **A**. Bubble graph of GSEA of TGAC8 SAN transcriptome. X-axis is normalized enrichment score (NES) in GSEA, bubble size represents number of genes in a term, and bubble color indicates enrichment using four different databases (KEGG, GO_CC, GO_MF, GO_BP). **B**. Two representative GSEA enrichment plots. Top: enriched using KEGG database; bottom: GO_BP.

In terms of cardiac function, we enriched apelin cardiomyocyte signaling pathway in IPA (Figure 3B). Apelin plays an important role in the regulation of cardiac functions, including inducing vasodilation, raising contractility and HR, and reducing reperfusion injury^18–20^. From the apelin cardiomyocyte signaling pathway generated in IPA, we observed that the transcriptional expression of myosin light chain genes was decreased in TGAC8 vs. WT, but the transcriptional expression of genes encoding several enzymes, including phosphatidylinositol-4,5-bisphosphate 3-kinase, phospholipase C, and protein kinase C, was increased in TGAC8 vs. WT (Figure 8A). To corroborate the enrichment of KEGG pathway, cardiac muscle contraction (Figure 4), we searched the enriched pathways within GSEA and found expression of most genes in the cardiac muscle contraction pathway was declined and the peak was downward in TGAC8 mouse (Figure 8B). The two enrichments in IPA and KEGG prompted us to further study the possibility of cardiac function impairment.

**Figure 8.**
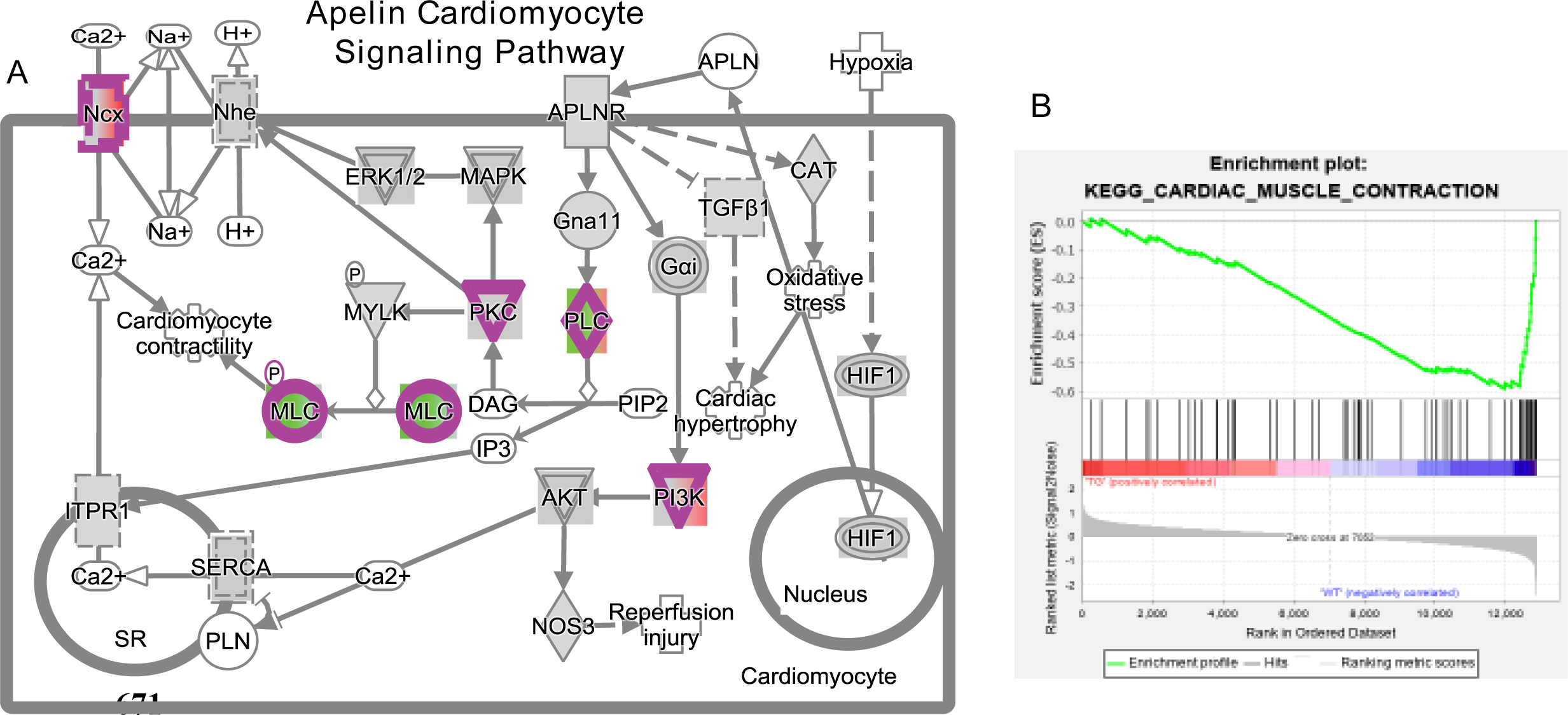
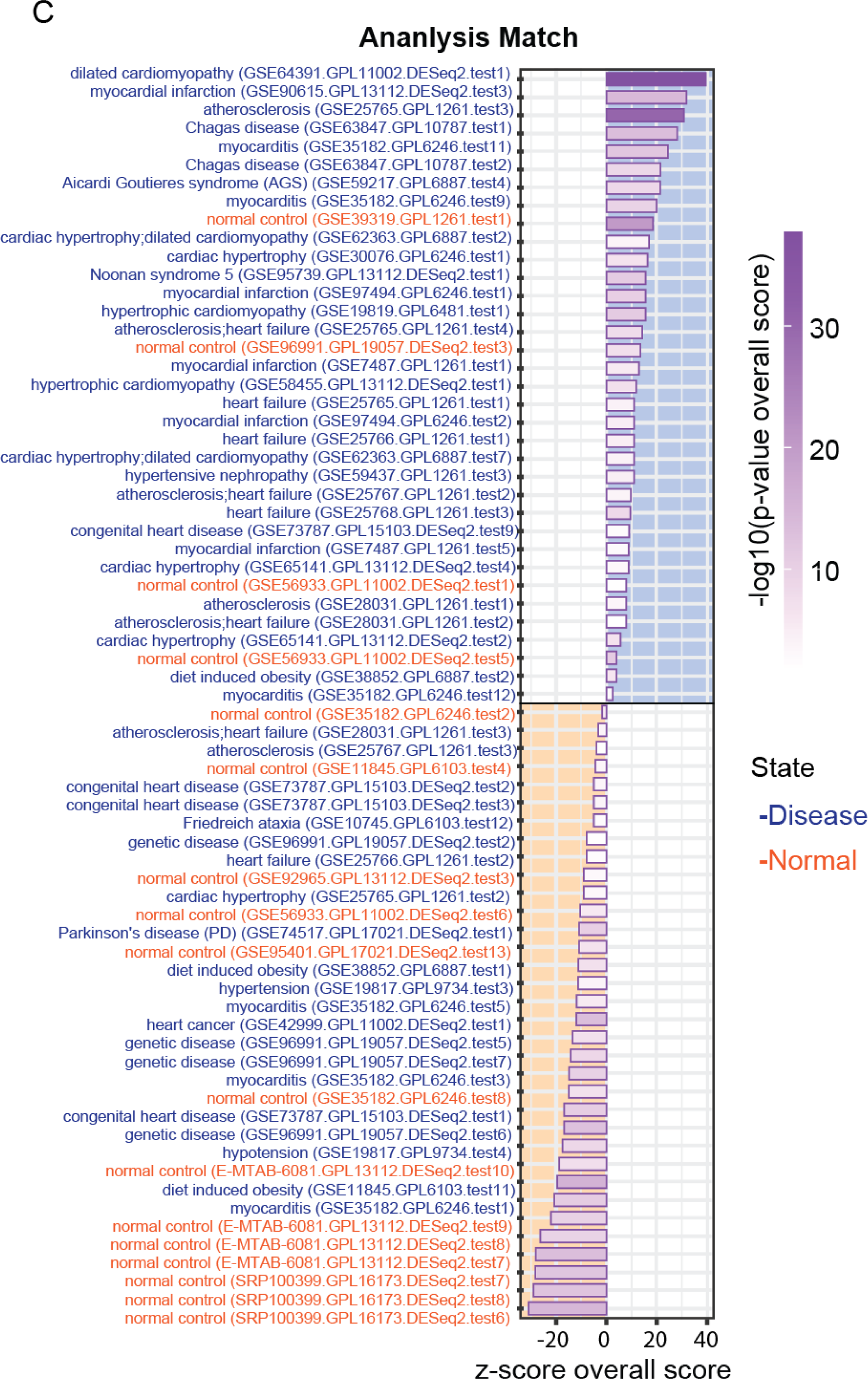
Association with impaired cardiac functions in TGAC8 SAN. **A**. Visualization of changed genes in the term of apelin cardiomyocyte signaling pathway, generated in IPA. Genes with purple outline are changed significantly. Green represents downregulated, and red represents upregulated. **B**. GSEA enrichment plot of cardiac muscle contraction. **C**. Bar plot of analysis match with public projects in IPA. X-axis is z-score overall score. Positive and negative scores indicate positive and negative correlations respectively. Color gradient represents -log10(p-value overall score), indicating the confidence of the analysis match. The darker, the high confidence. Y-axis is public project’s name. Blue indicates disease state, and red indicates normal state.

By the utilization of databases in IPA, we compared the TGAC8 mouse SAN transcriptome with all available public projects and displayed the analysis match in Figure 7C. Obviously, there were more matched disease projects with positive z-score, indicating that TGAC8 SAN transcriptome was more comparable to/like heart disease states than normal state (Figure 8C). Therefore, the reprogramed SAN transcriptome implied that the chronic overexpression of AC8 in the heart increases the HR at the cost of cardiac function and may result in/be associated with CVDs in long term.

### 6. Regulatory pathway of TGAC8 SAN transcriptome

To clarify the mechanistic pathway, we performed the prediction of possible upstream regulators mediating the effect of overexpression of AC8 to reprogram the SAN transcriptome. As RNA-seq data mainly reflects the regulation at transcriptional level, we first set the filter to screen all transcription regulators. Many predicted upstream regulators were also identified as significantly changed genes in the TGAC8 SAN transcriptome, so we set the second filter to select those genes whose transcriptional expression was changed in the same direction to the trend of predicted activation. Finally, we ranked all remained genes by B-H corrected p-value of prediction. Two regulators, PPARGC1A and KDM5A, were the most significantly inhibited and activated factors respectively. Peroxisome proliferator-activated receptor gamma coactivator 1-alpha (PGC-1α), encoded by the *PPARGC1A* gene^21^, interacts with many other regulators, including peroxisome proliferator-activated receptor gamma (PPAR-γ or PPARG), cAMP response element-binding protein (CREB), and nuclear respiratory factors (NRFs)^22–24^, to regulate mitochondrial biogenesis and energy metabolism^25^. Lysine-specific demethylase 5A (KDM5A) is encoded by the *KDM5A* gene^26^, and its loss restores mitochondrial function in cells lacking Rb1, partly by activating PGC-1α^27^. Therefore, we presumed the two transcription regulators as the mediators of the effect of overexpression of AC8 in SAN, and thus we established a regulatory pathway starting from the two regulators within IPA (Figure 9A). In the established pathway, the majority of genes under the control of the two regulators encode subunits in mitochondrial complex I, III, IV and V, whose downregulations contribute to mitochondrial dysfunction and disrupted OXPHOS process. In addition, the protein, Mitochondrial Ribosome Protein L53 (MRPL53), which is translated from one downstream gene whose expression is under the control of KDM5A, is a subunit of mitochondrial ribosome, and another protein, Translocase Of Outer Mitochondrial Membrane 7 (TOMM7), which is translated from another gene, is important in import of molecules into mitochondria. Therefore, the activated KDM5A could also suppress other mitochondrial functions such as mitochondrial translation and transport.

**Figure 9.**
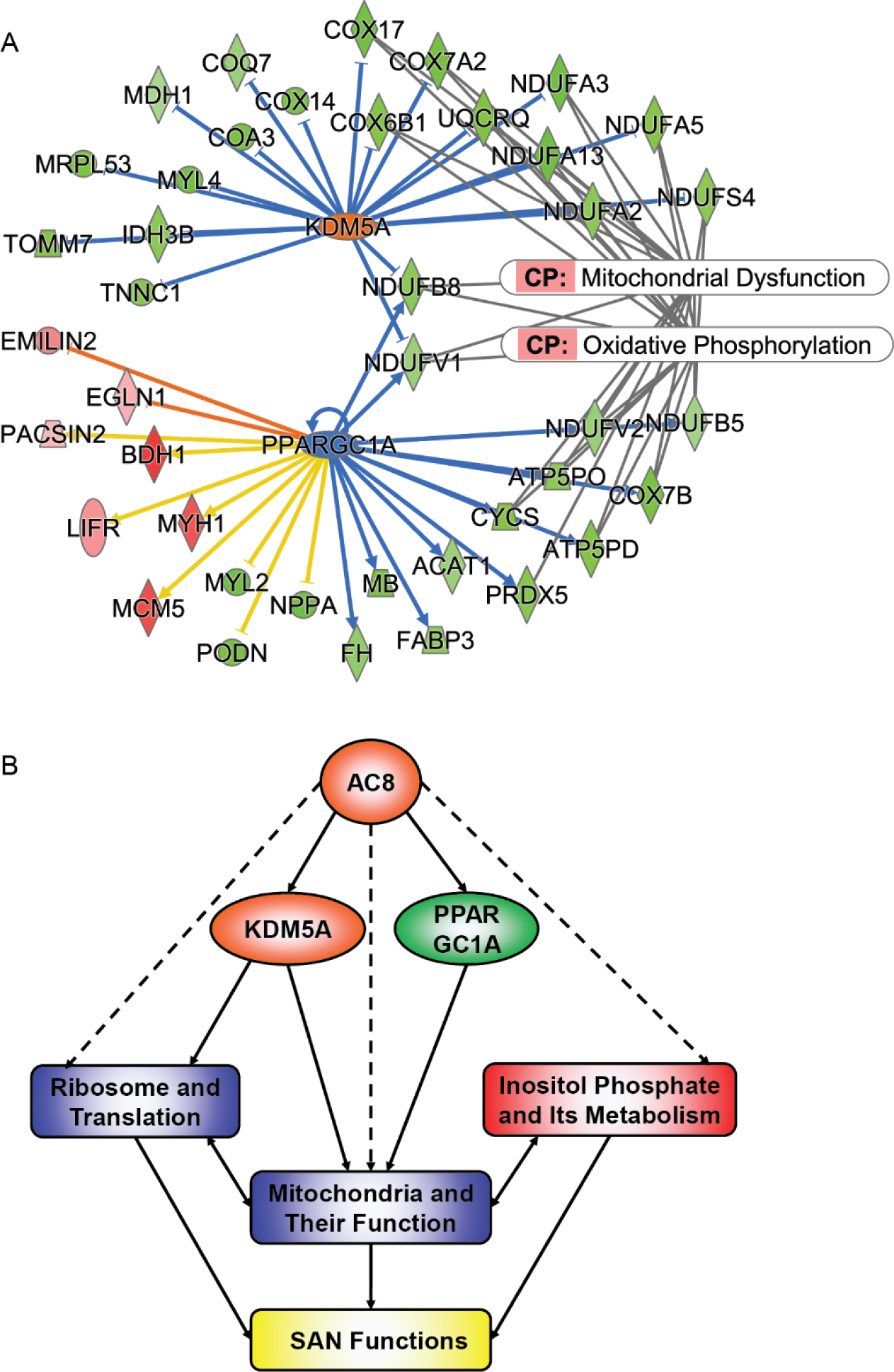
Regulatory pathway of TGAC8 SAN transcriptome. **A**. Mechanistic pathway from predicted upstream regulators to regulated downstream pathways. Two oval dots in the center denote the upstream regulators predicted on the basis of TGAC8 SAN transcriptome. Blue indicates inhibited, and orange indicates activated. Surrounding dots denote genes that are under the control of the two upstream regulators and changed significantly in TGAC8 SAN transcriptome. Green indicates downregulation, and red indicates upregulation. Edges between dots represent relationships between genes. Arrows denotes promotion, and short bar denotes suppression. Blue and orange edges indicate that the predicted activation state of upstream regulator leads to the gene downregulation and upregulation respectively in TGAC8 SAN transcriptome. Yellow edges represent findings inconsistent with state of downstream molecule. Two text boxes on the right represent downstream pathways that are enriched from differentially expressed genes. **B**. Deduced schematic: Transgenic overexpression of AC8 in SAN suppresses ribosome and translation, and mitochondria and their functions, while promotes inositol phosphate and its metabolism, thus associated with impaired SAN functions. Two transcription regulators, KDM5A and PPARGC1A, are possible mediators of overexpression of AC8.

## Discussion

In SAN, the ACs-cAMP-PKA signaling pathway plays an important role in pacemaking^3,5^. Previously, we discovered that the cardiac-specific overexpression of AC8 elevates the HR, but the overall pattern of marked coherency or loss of complexity (HR variability) within the TGAC8 heart rhythm is similar to that associated with aging or cardiac pathology^7^. In this research, we conducted RNA-seq to analyze the SAN transcriptome to reveal the underlying molecular mechanism and envisioned a possible long-term effect of chronic overexpression of AC8 in the heart. As expected, we observed obvious reprogramming in the SAN transcriptome (Figure 1-2). Beyond our expectation, it was mitochondrial function that was enriched most significantly and consistently in all analyses using multiple bioinformatic methods. We also enriched the suppressed ribosome and translation, and promoted inositol metabolism. Integrating all analyses, we deduced a schematic (Figure 9B). The cardiac-specific overexpression of AC8 suppresses ribosome and translation, and mitochondria and their functions, while promotes inositol phosphate and its metabolism. Two transcription regulators, KDM5A and PPARGC1A, may mediate the effect of overexpression of AC8 on ribosome and mitochondria. The abnormal ribosome and inositol phosphate could also cross talk with mitochondria to disrupt mitochondrial functions. In long term, the chronic overexpression of AC8 increases HR overtime at the cost of impaired SAN functions, perhaps resulting in/associated with CVDs.

### 1. Why does TGAC8 mouse SAN inhibit mitochondria and ribosome while activating inositol phosphate metabolism?

In our previous work, we observed the elevated HR in TGAC8 mouse relative to WT mouse^7^. The high-intensive and restless cardiac contraction lays a heavy load on energy production in mitochondria, which is the hub of energy homeostasis and metabolic homeostasis. The energy in shortage is hard to meet the demand for protein synthesis and cardiac contraction at the same time, so the translation could be sacrificed, as shown in the enrichments of suppressed ribosome components and translation in this research. The temporary or moderate decline of translation could induce several beneficial stress responses, including endoplasmic reticulum (ER) unfolded protein response (UPR) (UPR^er^) and mitochondrial UPR (UPR^mt^), to improve the protein quality control^28^. Interrogating the processed TGAC8 SAN RNA-seq data, we found some factors involved in UPR were upregulated, e.g., Atf6, Mbtps1, Mbtps2, Ern1, etc.

On the other hand, the regulated mitochondrial function is often accompanied with altered metabolism. Inositol phosphate, a type of metabolic derivative from lipids, is often localized in plasma membrane^29^, where it affects Ca^2+^ release from intracellular stores^30^ and regulates Ca^2+^ flow between mitochondria and endoplasmic/sarcoplasmic reticulum^16^. In the heart, the upregulated inositol phosphate increases the inner cellular Ca^2+^ to enhance cardiac contraction^14,15^, which may be an additional arm to PKA pathway in mediating the effect of activation of AC8 to increase the HR^7^. Therefore, TGAC8 mouse SAN prefers to inhibit mitochondria and ribosome while activate inositol phosphate metabolism to maintain high-intensive cardiac contraction.

### 2. How do the regulated pathways mediate the effect of TGAC8 in long term?

The three hub pathways, mitochondrial function (Figure 5), ribosome and translation (Figure 6), and inositol metabolism (Figure 7), were not distinct from each other. Mitochondria not only produce energy for protein synthesis, but also control metabolism to impact inositol phosphate indirectly. In turn, the assembly of complete functional mitochondrial complex relies on proteins synthesized in cytosol and mitochondria. IP3 regulates mitochondrial Ca^2+^ uptake to influence ATP production^17^. Although the temporary or moderate protein stress and mitochondrial stress are beneficial to health, the long-term and vigorous stress is detrimental to health^31^. For example, the elevated HR increases blood supply in response to exercise or emergency, but it aggravates the burden of heart in the context of chronic heart diseases. Similarly, the abnormally and persistently activated IP3Rs signaling may result in arrhythmias, hypertrophy and even heart failure^14,15^. Under chronic CVDs states, UPR may be maladaptive^32^, exemplified by its arrhythmic effect during human heart failure by affecting cardiac ion channels expression^33^. The persistent UPR could also be an upstream signal to promote cancer^34^. In our TGAC8 mouse model, we observed elevated HR as a chronic stress laid on mouse heart^7^. The enriched cardiac function pathways, especially downregulated cardiac contraction pathway, predict a coming cardiac disorder, in line with a recently published work where aggravated age-related myocardial dysfunction was reported in TGAC8 mice compared with their WT littermates^35^. Therefore, our results not only support the concept of detrimental role of chronic stress, but also indicate the association of early-stage reprogramming of SAN transcriptome with the late-stage cardiac pathology.

### 3. Can we regulate these pathways to ameliorate the side-effect of TGAC8 in long term?

As the robust functions of ribosome and mitochondria are pivotal to health, the ideal way to improve cardiac fitness is to concurrently enhance cardiac contraction and biological processes, i.e., OXPHOS, translation, and metabolism, to maintain cardiac homeostasis at a higher level. The TGAC8 has already elevated the HR, so we need to increase the functions of ribosome and mitochondria to maintain health. Two transcription regulators, KDM5A and PPARGC1A (PGC-1α), were predicted to be the mediators in TGAC8 to inhibit ribosome and mitochondria. The expression of KDM5A was reported to be negatively correlated with subunits in mitochondrial complexes^27^, and PGC-1α is well known as a positive regulator of mitochondrial biogenesis^25^. Therefore, the two regulators have the potential to be therapeutic targets in improving SAN functions, particularly in aged TGAC8 mouse, or patients under similar chronic stresses.

### 4. Limitation

We unraveled the reprogramed transcriptome in TGAC8 vs. WT SAN, which indicates the suppressed components and function of ribosome and mitochondria, along with promoted inositol phosphate metabolism, but more wet lab experiments were not included in this research due to the scarcity of isolated SAN samples. In addition, the complete and clear signaling pathway, from AC8 to mediators, to target genes, and to downstream effects, need further validation. For example, the double gene mutation mouse models, e.g., TG^AC8^-TG^PPARGC1A^ or TG^AC8^-KO^KDM5A^ mouse, could be used to in-depth study the regulatory mechanisms of overexpression of AC8 in SAN and to reveal potential therapeutic targets of chronic CVDs, especially during aging.

## Acknowledgements

We thank all members in Laboratory of Cardiovascular Science, National Institute on Aging, National Institutes of Health, for technical assistance and discussion. This work utilized the computational resources of the NIH HPC Biowulf cluster (http://hpc.nih.gov).

